# A consortium of human commensals protects against middle ear colonization by otopathogens

**DOI:** 10.64898/2026.01.26.701666

**Authors:** Kalyan K. Dewan, Maiya Callender, Jillian Masters, Emily A. Gilbertson, Jillian Hurst, Eric T. Harvill

**Affiliations:** Department of Infectious Diseases, College of Veterinary Medicine, University of Georgia, Athens, Georgia, Untied States; Division of Infectious Diseases, Duke University School of Medicine, North Carolina, United States

## Abstract

A year-long sequencing analysis of bacterial commensals sampled from infants during periods in which they were healthy or suffering recurrent ear infections [otitis media (OM)] identified several species of bacterial commensals that correlate with health and absence of ear infections. Here we consider and test the possibility of a causal relationship between a group of commensals and periods of health. We assemble a set of five health-associated bacterial species into a nasopharyngeal commensal consortium (NPCC) and test whether these organisms can effectively colonize the respiratory tracts of mice so that their effects on invading pathogens could be evaluated. We observed that NPCC efficiently colonize mice and that they provide substantial protection against the otopathogens, *Streptococcus pneumoniae* and *Bordetella pertussis*, reducing numbers of each in the middle ears by 99 to 99.9%. The NPCC also affected colonization/growth of these pathogens within the lower respiratory tract, suggesting complexity in these interactions. Together these data demonstrate a profound effect of commensals on invading otopathogens and describe a powerful experimental system in which the important interactions between the healthy infant microbiota and invading pathogens can be studied mechanistically.

## INTRODUCTION

The vast majority of children suffer from ear infections (otitis media, OM) [1], with considerable diversity in presentation and outcomes that range from minor and transient self-limiting ailments to chronic or extreme and debilitating infections requiring multiple rounds of antibiotics and surgery. Over the past decades a broad range of factors (genetic and environmental) have been associated with the likelihood of individuals to suffer from respiratory and middle ear diseases and to the sequelae of these infections [2]. More recently, growing numbers of clinical studies across the globe [3–11] have been reporting correlations between ear infection occurrence and the populations of microorganisms residing in the mucosal surfaces of the respiratory tract (and adenoids), collectively referred to as the respiratory microbiota, with the propensity of individuals to suffer infections or remain healthy [12–15].

It would be reasonable to speculate that resident microbiota of the respiratory tract impact colonization by otopathogens [16]. Invading pathogens must pass through the complex sol-gel mucus layers that are already colonized by the diverse array of commensal microorganisms that constitute the respiratory microbiota [17]. Individual microbes within this mucoidal layer are in stiff and continual competition with other residents and potentially with organisms/pathogens inhaled with each breath. Such competition can be direct via bacteriocins [18–20] and other selectively toxic molecules [21] secreted from dedicated systems [22, 23], or can be indirect, involving depletion of scant but necessary nutrients in the shared environment [24, 25]. Alternatively, competition may be “apparent”, for example as when mediated by phage that preferentially predate the invader, or induction of host responses that act differentially on the resident and invader [26, 27]. As described for other anatomic sites [28, 29], populations of microorganisms can establish relatively stable consortia of microbes that persist together over time, collectively repelling the near constant barrage of potential invaders. Yet, unlike the microbiota of healthy adults where the diverse communities of commensals appear stably established in equilibrium with each other and the host immune system, the microbiota in newborns undergo a succession of compositional changes over the early years of an infant’s life that, maturing along with the neonatal-infant immune system. This period of transitional microbiome correlates with the period of high susceptibility to OM [30, 31] further implicating the resident microbiota in affecting susceptibility/resistance of the middle ear to invading pathogens.

In a longitudinal study examining the microbiota of infants who suffered recurrent ear infections (n= 58, followed for 1-year), [32], we identified distinct commensal taxa that expanded in population size during periods of health compared to periods of infection. Identification of these species was further supported by analysis of previously published cohorts that described differences in respiratory microbiome composition among healthy children and those with viral and bacterial respiratory infections [5, 33–36], as well as data from in vitro experiments. These observations establish that alternate sets of bacterial commensals are associated with the “infection-free” versus the “infected” status of susceptible infants, and raise the compelling hypothesis that specific commensal bacteria may be protective against ear infections.

Here, to more directly test for a causal relationship, we have assembled a consortium comprising equal numbers of five bacterial species that were associated with periods of health, which we refer to as the Nasopharyngeal Commensal Consortium (NPCC). We then tested the ability of the NPCC to colonize the mouse respiratory tract, and examined the potential protective effects that the NPCC might have against colonization of the mouse middle ears and respiratory tract by two human pathogens, the gram-positive *Streptococcus pneumoniae* (*Spn*) and gram-negative *Bordetella pertussis* (*Bp*). Both pathogens target infants causing serious respiratory infections and are also associated with otitis media. The NPCC successfully colonized the respiratory tract and spread amongst neonates, protecting their ME against both pathogens. They had more nuanced effects in adult mice. Together these results reveal substantial effects of commensals on invading pathogens and define an experimental system that is well suited to probe the complex intermicrobial competition of the infant respiratory microbiota and invading otopathogens and how these differ in adults and infants.

## MATERIALS AND METHODS

### Bacterial Strains and Growth

Growth and maintenance of all bacterial species used in this study was conducted in the BSL-2 certified Harvill lab at the University of Georgia following procedures approved by the IBC, University of Georgia.

#### Nasopharyngeal commensal consortium (NPCC)

Commensal bacteria of the NPCC used in this study (*Moraxella nonliquefaciens*, *Streptococcus mitis*, *Corneybacterium accolens*, *Corneybacterium propinquum*, *Dolosigranulum pigrum*) (*Supplementary Figure 1*) were purchased from American Type Culture Collection (ATCC; Manassas VA). Liquid cultures of each species were grown in Brain Heart Infusion (BHI) (Becton Dickenson) broth. BHI was supplemented with 0.1% Tween-80 for *Corynebacterium* spp. or 60 mM MOPS for *D. pigrum*. CFU of individual cultures enumerated by plating on Tryptic Soy agar with 5% defibrinated sheep blood. The inoculum of all five bacterial species (NPCC) was prepared by mixing an estimated 1×10^7^ CFU/mL of each of the 5 species in PBS (5×10^7^ CFU/mL of culturable bacteria). Aliquots of inoculation stocks were frozen with 10% glycerol and stored at - 80^0^C. Bacterial numbers were enumerated to confirm the delivered CFU numbers for each inoculation.

*Streptococcus pneumoniae* TIGR4 used in this study was kindly provided by Prof Balazs Rada from his laboratory stocks at the University of Georgia. For enumerating CFU, bacteria were plated on Tryptic Soy agar (BD) with 5% defibrinated sheep blood (Hemostat Laboratories) supplemented with 4 μg/mL of gentamicin as a selective agent. Liquid cultures of the bacteria for inoculations were prepared using BHI (BD) grown overnight at 37°C with shaking (200 rpm). Bacterial numbers were estimated spectrophotometrically at OD of 600 nm (0.1 ≅ 2 x10^7^ CFU/mL) and inoculation stocks at required concentrations prepared by diluting in PBS.

*Bordetella pertussis* Tohama I (gentamicin resistant derivative) used in this study was from the Harvill laboratory stocks. Bacterial CFUs were enumerated by plating on Bordet-Gengou agar (DIFCO) supplemented with 10% sheep blood (Hemostat laboratories) and 20 mg/mL gentamicin (Sigma), and incubated for 4-5 days at 37°C. Liquid cultures of the bacteria for inoculations were grown in Stainer-Scholte broth (with 20μg/mL gentamicin) at 37°C with shaking (200 rpm). Bacterial numbers were estimated spectrophotometrically at OD of 600 nm (0.1 ≅ 2 x10^8^ CFU/mL) and inoculation stocks at required concentrations prepared by diluting in PBS.

### Mouse Experiments

Six- to eight-week-old female and male C57BL/6J, were procured from the Jackson Laboratory (Bar Harbor, ME) and maintained for experiments that employed adult mice (6-8 weeks old, n= 4 -5 mice/group) and for breeding (2-6 months old) to generate pups (8-10 pups/litter group) used in this study. All mice were maintained in specific pathogen-free facilities at the animal facilities at the University of Georgia and procedures conducted following approved institutional animal use guidelines.

#### NPCC colonization

For colonizing pups with the NPCC, 2-day-old pups (P2) were lightly sedated with 5% isoflurane (Pivetal) and inoculated by droplet inhalation of ∼ 2.5 x 10^4^ CFU of the NPCC delivered in 2.5 μl of PBS. For experiments involving colonization of adults, the mice were inoculated with ∼ 5x 10^5^ CFU delivered in 10μL of PBS. Inoculated adults and pup litters were left for 3 days for NPCC to establish colonization.

#### Pathogen challenge

Lightly sedated 5-day old (P5) pups were inoculated by droplet inhalation of ∼10^4^ CFU of either *S. pneumoniae* or *B. pertussis* resuspended in 15μL of PBS delivering the pathogen across the respiratory tract. For adults, mice were inoculated with ∼ 2 x10^5^ CFU of the pathogens delivered intranasally in 10μL of PBS, limiting the initial challenge to the upper respiratory tract. Bacterial loads across the organs for *S. pneumoniae* inoculated pups and adults were assessed after 24 hours, while *B. pertussis* inoculated pups and adults were assessed after 72 hours.

#### Euthanasia and organ harvesting

At the indicated timepoints, mice (adults and pups) were euthanized via CO_2_ inhalation (4L min^-1^) over 2-3 minutes. Cervical dislocation was performed for all adult mice post CO_2_ exposure as secondary procedure to ensure full euthanasia. For infant mice, the secondary procedure was decapitation post C0_2_ exposure. The nasal cavities (adults)/nasopharynx (infant mice), lungs and middle ears of euthanized mice were carefully excised and the organs were homogenized in a bead mill beater in 1 mL of cold PBS. CFU numbers were enumerated by plating aliquots of the homogenized samples on appropriate agar and supplements to quantify culturable bacteria. Where 16S sequence analysis of microbiota was performed, the remaining homogenized tissue was centrifuged (12,000 rpm for 5 minutes), supernatant discarded, and pellets stored at -80^0^C for further processing.

#### 16S rRNA gene amplicon sequencing and analysis

The samples were processed and analyzed with the ZymoBIOMICS^®^ Targeted Sequencing Service (Zymo Research, Irvine, CA). The ZymoBIOMICS^®^-96 MagBead DNA Kit (Zymo Research, Irvine, CA) was used to extract DNA using an automated platform. The library of whole 16S sequencing was prepared by following the full-length 16S amplification protocol from PacBio [**37**]. In brief, the whole 16S gene was amplified using the 27F (AGRGTTYGATYMTGGCTCAG) and 1492R (RGYTACCTTGTTACGACTT) primers with barcodes and adapters. Two ng of DNA was used as the PCR template for each sample, and the PCR was run under the following conditions: initial denaturation at 95 °C for 3 minutes, followed by 25 cycles of 95 °C for 30 seconds, 57 °C for 30 seconds, 72 °C for 60 seconds. PCR products were subjected to clean up with Select-a-Size DNA Clean & Concentrator MagBead Kit (Zymo Research, Irvine, CA) to retain fragments >300bp. Each reaction library was quantified by NanoDrop and pooled together with equal DNA mass. Pooled libraries were prepared for sequencing using the SMRTbell^®^ prep kit 3.0 (PacBio, Menlo Park, CA). Control samples were prepared in parallel with the experimental samples, including the ZymoBIOMICS^®^ Microbial Community Standard (Zymo Research, Irvine, CA) as a positive control for DNA extraction and library preparation. Negative controls included blank extraction controls and blank library preparation controls. Libraries were sequenced on one 8M SMRT cell on the Sequel IIe system (PacBio, Menlo Park, CA). Sequencing reads were analyzed using the DADA2 pipeline [38], including removal of potential sequencing errors and chimeric sequences. Taxonomy was assigned using Greengenes2 [39]. Presumed reagent contaminant ASVs were removed using the frequency method in the *decontam* R package version 3.20 (threshold = 0.10) [40]. The median (IQR) reads per sample were 13,640 (5,409–33,419) after quality filtering. There were 3,206 ASVs mapping to 190 genera and 200 species.

#### Microbiome analysis

ASVs were filtered to remove low-abundance sequences with ≤10 total counts across all samples to reduce noise from potential sequencing artifacts. Known environmental contaminants were identified based on literature and removed from the dataset, including genera commonly associated with laboratory or reagent contamination (*Jeotgalicoccus, Agrobacterium, Bradyrhizobium, Sphingopyxis, Herbaspirillum, Rubrivivax, Pedomicrobium, Leifsonia, and Methyloversatilis*). We applied centered log-ratio (CLR) transformation after adding a pseudocount of 1 to handle zero values. For CLR transformation, samples were first converted to relative abundances, then the geometric mean of each sample was calculated, and each taxon abundance was log-transformed and centered by subtracting the geometric mean. To determine successful engraftment of NPCC member species, we extracted read counts for target taxa Multiple ASVs representing the same species were combined by summing their read counts. Relative abundances were calculated as percentages of total reads per sample, and mean relative abundances were computed for each treatment group. Overall microbiome composition was visualized using stacked bar plots showing mean relative abundances of the 30 most abundant taxa, calculated from non-transformed count data. Taxa were aggregated at the genus-species level by combining ASVs with identical taxonomic assignments. The “Other” category represented all remaining taxa. To assess whether NPCC inoculation significantly altered overall community composition, we performed principal coordinates analysis (PCoA) based on Euclidean distances calculated from CLR-transformed abundances. Statistical significance of compositional differences among treatment groups was tested using permutational multivariate analysis of variance (PERMANOVA) with 999 permutations, implemented via the adonis2 function in the vegan package. To identify specific taxa that changed in abundance following NPCC inoculation, we performed differential abundance analysis comparing control and treated groups for each tissue type. Linear models were fitted for each ASV using CLR-transformed abundances as the response variable and treatment group as the predictor. To control for multiple hypothesis testing across all ASVs, we applied the Benjamini-Hochberg false discovery rate (FDR) correction. Taxa were considered significantly differentially abundant at an FDR-adjusted p-value < 0.05. We also reported nominally significant taxa (unadjusted p < 0.05) to identify potential biological signals that may require validation in larger studies. All statistical analyses were performed in R version 4.4.2. Data manipulation utilized *dplyr* and *tidyr* from the *tidyverse* package suite. Visualizations were created using *ggplot2* with color palettes from *RColorBrewer*. Multi-panel figures were assembled using *cowplot*. All statistical tests were two-sided, and significance was defined as p < 0.05 unless otherwise specified.

#### Statistics

Data generated for CFU enumeration was evaluated using the Students t-test from Graphpad prism (V2.0).

### Ethics Statement

This study was carried out in accordance with the recommendations in the Guide for the Care and Use of Laboratory Animals of the National Institutes of Health. The protocols followed was approved by the Institutional Animal Care and Use Committees at The University of Georgia at Athens, GA (AUPs: Bordetella-Host Interactions: A2025 04-002-Y1-A1, Breeding Protocol: A2025 04-006-Y1-A1, Maternal vaccinations in neonatal protection: A2024 06-019-Y2-A2, and Neonatal models of Bordetella infection 2.0: A2025 06-027-Y1-A1). Individual experimental mice were monitored daily for signs of distress over the course of the experiments to determine if they required to be euthanized to prevent unnecessary suffering.

## RESULTS

### 1. Members of the NPCC colonize the infant mouse nasopharynx

To examine if the NPCC would colonize the infant nasopharynx and how this might impact the microbiota, 5-day old (P5) pups were inoculated with a size-adjusted dose of ∼2.5 x 10^4^ CFU of the NPCC in a volume of 2.5 μL of PBS. A second litter of P5 pups from the same breeding colony was left untreated to serve as controls. 3 days later pups were euthanized and nasopharynx, MEs and lungs processed for analysis.

#### Culturable bacterial load

We first assessed how treatment of infant mice with the NPCC would affect the culturable bacteria in various organs. **Figure 1** compares the bacterial CFU recovered on blood base agar from homogenates of the nasal cavities (NC), middle ears (ME) and lungs (LNG) of the untreated (black columns) and NPCC-treated (grey columns) infant mice. While untreated pups showed relatively low numbers of bacteria in their NCs (∼10^4^ CFU), MEs (∼10^3^ CFU) and LNGs (10^1^ CFU), three days following the NPCC inoculation the number of bacteria increased from 10 to 1000-fold across the three organs (NC ∼ 10^5^ CFU, ME ∼ 5 x10^3^ CFU, LNGs ∼10^3^ CFU). These observations indicate that the administration of NPCC markedly alters the distribution of microbiota across these organs.

**Figure 1.**
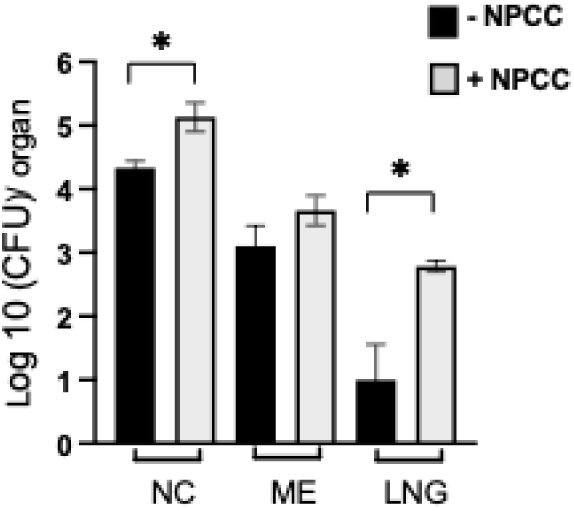
Bacterial recovery across organs of infant mice following NPCC administration. Graph shows CFU numbers of culturable bacteria recovered from the nasal cavities (NC), middle ears (ME) and lungs (LNG) of P8 pups that were either untreated (black columns) or inoculated 3 days earlier (at P5) with NPCC (grey columns). Error bars: S.E.M; *=p<0.05. (Results compiled from two experiments)

#### NPCC colonization and effect on respiratory microbiota in pups

To examine the relative abundance of the NPCC member species, we conducted long-read 16S rRNA DNA sequencing to identify the bacterial compositions in the 3 dpi organ homogenates of NPCC-treated and untreated pups. We first evaluated the mean relative abundance of the five NPCC component species (**Figure 2**). We did not detect amplicon sequence variants (ASVs) assigned to any of these species in tissues from untreated animals. Notably, we did not identify ASVs corresponding to *M. nonliquefaciens* or *C. propinquum* in any of our samples; however, all other component species were detected in 4 of the 5 nasal samples. We found that *C. accolens*, *D. pigrum*, and *S. mitis* made up approximately 1% of detected species in the nasal cavity, while *S. mitis* and *D. pigrum* made up less than 0.05% of taxa identified in the middle ear. NPCC species were not detected in lung samples.

**Figure 2.**
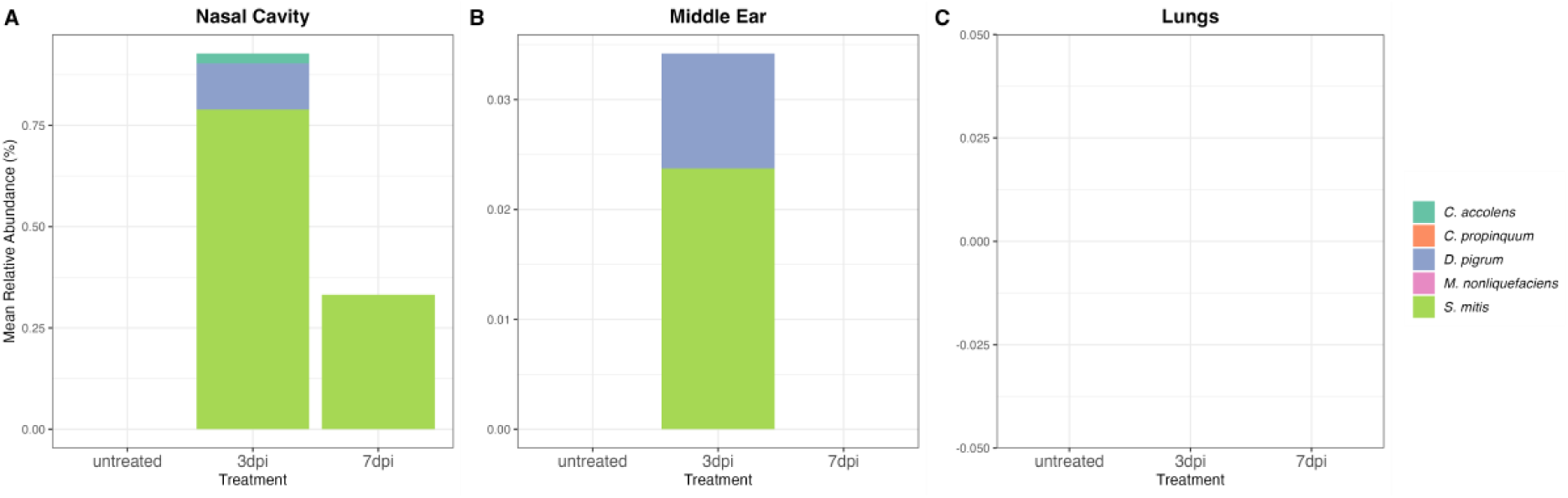
Mean relative abundance of NPCC species in the respiratory tracts of infant mice. Long-read 16S rRNA sequencing was used to profile bacterial species in respiratory tissues collected from infant mice 3 and 7 days after inoculation (3dpi, 7dpi) after inoculation with the NPCC. Graphs show the mean relative abundance of NPCC component species in A) nasal cavity tissues, B) middle ear tissues, and C) lung tissues from treated (n=5) and untreated (n=4) mice. NPCC species were not detected in untreated mice.

These results show that 3 of the 5 members of the NPCC had integrated into the nasopharyngeal microbiota at differing abundance levels (*S. mitis*> *D. pigrum*> *C. accolens*), with 2 species (*S. mitis> D. pigrum*) also observed in the MEs. We did not detect any NPCC species in lungs, indicating that the NPCC were not substantial components of the culturable bacteria in the lungs observed above (Figure 1).

#### NPPC impact on infant microbiota

To evaluate the impact of NPCC on microbiota composition in respiratory tissues we evaluated the top 30 taxa in organ homogenates from untreated and NPCC-treated pups 3dpi (Figure 3, Supplemental Figure 2). We found that NPCC treatment significantly altered microbial composition in the nasal cavity (PERMANOVA, p<0.0001), middle ears (PERMANOVA, p=0.004), and lungs (PERMANOVA, p<0.0001) on day 3 and 7 post inoculation when compared to untreated pups. Given that we observed significant compositional differences across all tissue types, we sought to identify individual taxa that were significantly altered between NPCC-treated and untreated animals. Using ordinary least squares linear regression, we identified 22 taxa that exhibited nominally significant changes between untreated and NPCC-treated animals; however, only a single taxon, *Limosilactobacillus reuteri* (ASV38), was significant after adjustment for multiple comparisons (estimate: 2.97, p_adj_=0.001). In the middle ears, we identified 2 taxa that exhibited nominally significant changes associated with NPCC inoculation (ASV9 and ASV11, both of which were classified as *Staphylococcus saprophyticus*), but neither was significant after adjustment for multiple comparisons. In lungs, we similarly identified four nominally significant taxa that did not survive adjustment for multiple comparisons (ASV3/*S. saprophyticus*; ASV1/*Streptococcus danielae*; ASV4/*S. danielae*; ASV134, unknown species).

**Figure 3.**
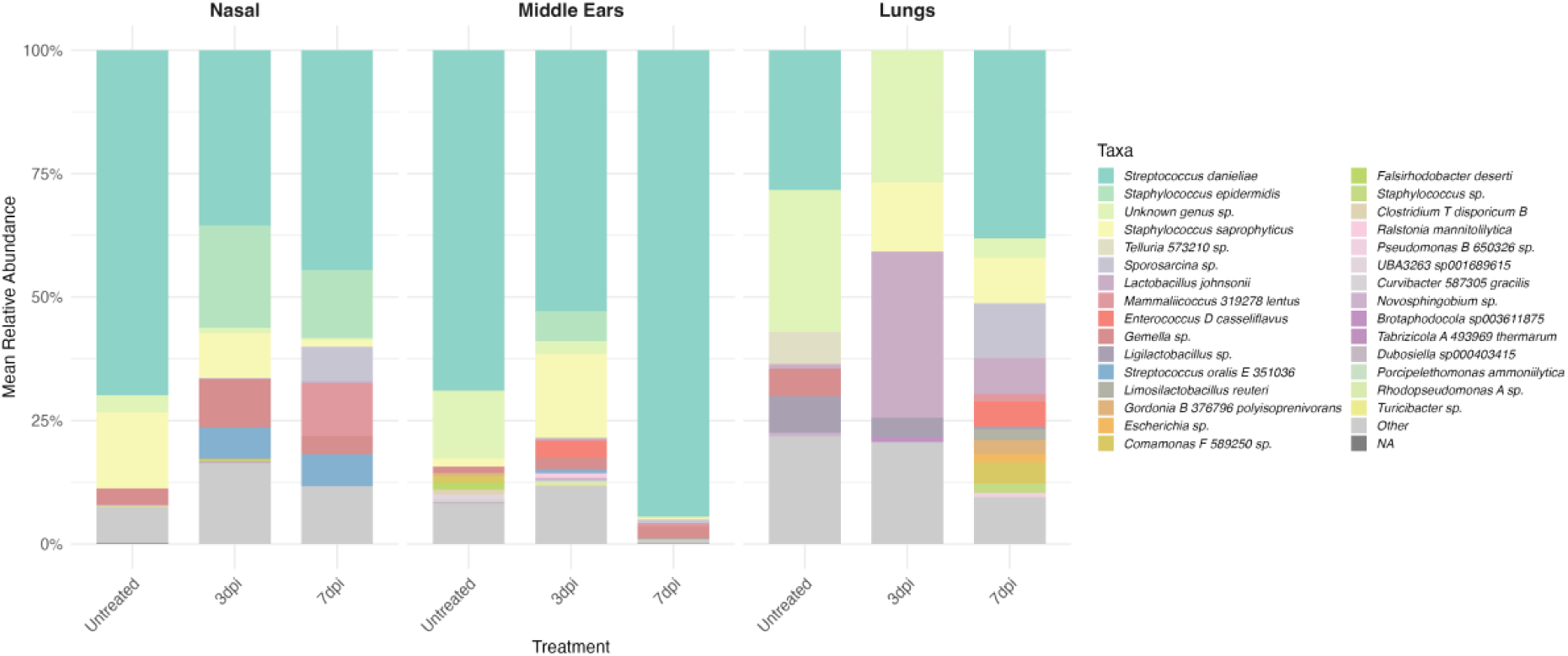
Impact of NPCC inoculation on microbiome composition of the respiratory tract. CLR-transformed abundances of the top 30 taxa are shown for nasal, middle ear, and lung tissue in untreated and treated animals 3 and 7-days post inoculation (dpi) of the NPCC.

**Figure 4.**
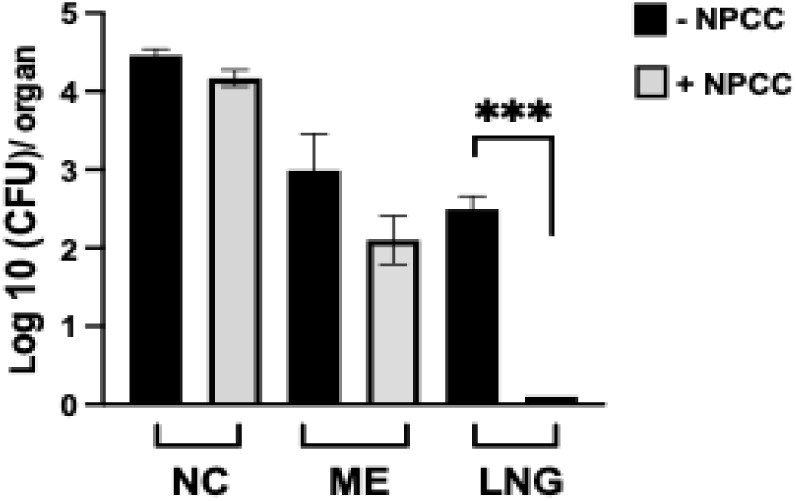
Effect of NPCC on *S. pneumoniae* colonization of infant mice. Graph shows CFU numbers of *S. pneumoniae* recovered from the nasal cavities (NC), Middle ears (ME) and lungs (LNG) of P9 pups, one day after being challenged. The pups were from litters that were either untreated (black columns) or inoculated with NPCC at P5 (grey columns). Error bars: S.E.M; ***=p<0.0001. (Results compiled from two experiments)

These results demonstrate that introduction of the NPCC induces complex readjustments of the bacterial populations across the organs of the infant mice. Interestingly, NPCC introduction was associated with significant expansion of *Lactobacillus johnsonni* in the lungs [41–43] and *Limosilactobacillus reuteri* in the nasal cavity [44–46]. Both species are being actively investigated for their beneficial immunomodulatory activities in the lungs and gut respectively. These observations indicate that aside from direct competitive advantages that the NPCC may bring against pathogens, more subtle advantages may be conferred likely via their competitive effects in reshaping the microbiomes of the respiratory tract.

### 2. NPCC impedes colonization of MEs by *Spn* and *Bp* in infant mice

Based on the compelling association between healthy microbes and resistance to OM, we tested the hypothesis that NPCC might protect infant mice from being colonized by human pathogens. We chose the gram-positive otopathogen *Streptococcus pneumoniae*, strain TIGR4, an invasive and well-studied pneumococcal strain that causes AOM in humans and is often used as a model pathogen to study AOM in mice. In addition, we also examined the effect of NPCC on *Bordetella pertussis*, a gram-negative pathogen and the etiological agent of pertussis (whooping cough), that primarily causes respiratory infections but has also been reported to ascend the Eustachian tube to infect the middle ear, causing OM among infants [47] and adult humans [48] and which we have noted in the middle ears of adult and infant mouse models of infection (unpublished observations). Litters of 5-day old (P5) C57Bl/6J pups were inoculated with 2.5×10^4^ CFU of the NPCC delivered in 2.5 μL of PBS, or left untreated. 3 days following NPCC administration the now 8-day old pups (P8) were intranasally challenged with an “infant-dose” of 1 x10^4^ CFU of either *S. pneumoniae* TIGR4, or *B. pertussis,* delivered in 15μL of PBS, a volume expected to reach the lungs of the inoculated pups (PMID: 39475256).

#### S. pneumoniae

The rapidly growing *S. pneumoniae* TIGR4 (doubling time:16 minutes; [49]) was analyzed one day post challenge (dpc) by plating dilutions from tissue homogenates on TSA plates (+5% blood) supplemented with 4μg/mL gentamicin, a concentration we found that was bactericidal to the NPCC and culturable native resident microbiota, but to which TIGR4 is resistant. Untreated pups (black columns) had large numbers of *S. pneumoniae* in the nasal cavity (∼4.5 x10^4^ CFU), and lower numbers in the lungs (∼500 CFU). All the pups not treated with the NPCC (-NPCC) also showed the MEs being colonized by S. pneumoniae to varying degrees (∼10^2^ - 10^3^ CFU) indicating that the pathogen had efficiently spread up the Eustachian tube to colonize the ME. By comparison, the +NPCC treated group (light gray columns) showed a modestly lower (∼0.5 log) numbers of *S. pneumoniae* in the NCs, and substantially lower numbers (∼10-fold) in the MEs, compared to untreated.

Interestingly, *S. pneumoniae* colonization of the lungs for the NPCC-treated group fell below the limit of detection (10 CFU) indicating the NPCC prevented *S. pneumoniae* reaching and colonizing the lungs.

#### B. pertussis

To assess the impact of the NPCC on *B. pertussis* colonization/growth, NPCC-treated and untreated pups were challenged with the pathogen as above (∼10^4^ CFU/15μL PBS). To allow sufficient time for the slower growth of *B. pertussis* (doubling time: 4-7 hours) colonization levels were assessed three days following this challenge (at P11). Untreated (-NPCC) mice showed high *B. pertussis* loads across the NC (∼10^5^ CFU), MEs (∼5 x10^3^ CFU) and LNGs (∼10^6^ CFU) (Fig.5), consistent with what we have observed earlier [50]. Interestingly *B. pertussis* loads of neither the NC nor LNG were affected by NPCC, but the *B. pertussis* numbers in the MEs were ∼10-fold lower (∼5×10^2^ CFU) in NPCC+ than in NPCC-pups, indicating NPCC provided substantial protection of the ME.

**Figure 5.**
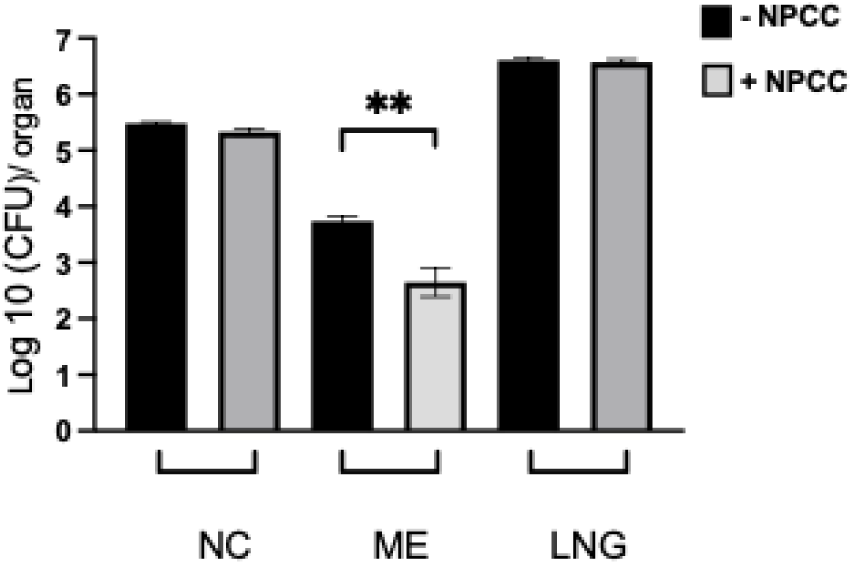
Effect of NPCC on *B. pertussis* colonization of infant mice. Graph shows CFU numbers of *B. pertussis* recovered from the nasal cavities (NC), Middle ears (ME) and lungs (LNG) of pups (P11), 3 days after being challenged. The pups were from litters that were either untreated (black columns) or treated with NPCC at P5 (grey columns). Error bars: S.E.M; **=p<0.001. (Experiment compiled from 2 experiments)

These results, showing measurable reductions of two highly diverse respiratory pathogens in the MEs of NPCC-treated infant mice, are in overall agreement with the associations of these commensals with a healthy middle ear status in human infants. These observations also indicate that this model can be used to investigate the mechanistic aspects of NPCC-mediated protection while more closely examining the different impact it appears to have on the colonization of lower respiratory tract.

### 3. NPCC impedes colonization of MEs by *S. pneumoniae* and *B. pertussis* in adult mice

We next examined if a similar ME protective effect of the NPCC would be observed in adult mice. To generate adult mice with or without the NPCC, groups of C57Bl/6J mice (6-8 weeks old) were either left untreated (controls) or inoculated with the adult dose of the NPCC (∼10^5^ CFU/10μL in PBS), as above. Three days later, both groups were challenged with ∼ 5 x10^5^ CFU of either *S. pneumoniae*, or *B. pertussis.* The inoculum was delivered in a relatively small volume of 10μL of PBS to localize the inoculum to the nasopharynx which would allow us to examine how the NPCC impacts growth and spread of the pathogens across the MEs and lower respiratory tract.

#### i) S. pneumoniae

Untreated (-NPCC) mice (black bars) had high numbers of *S. pneumoniae* (10^4^ to10^5^ CFU) in nasal cavities, middle ears and lungs, demonstrating that in the absence of NPCC *S. pneumoniae* efficiently colonizes, spreads and grows in the lungs and middle ears of adult mice (Figure 6). In stark contrast, mice previously treated with NPCC (grey bars) contained on average >95% fewer *S. pneumoniae*. The effect of NPCC was large but variable in the nasal cavity, and so did not show statistical significance, but differences were consistent, robust and significant in both middle ears and lungs, indicating that the NPCC substantially interfered with *S. pneumoniae* colonization/growth in these deeper organs. These results demonstrate that the NPCC impedes pathogen colonization/growth in complex ways and that the conditions of these infection assays provide an opportunity to study how commensals interfere with, and protect against, invading pathogens.

**Figure 6.**
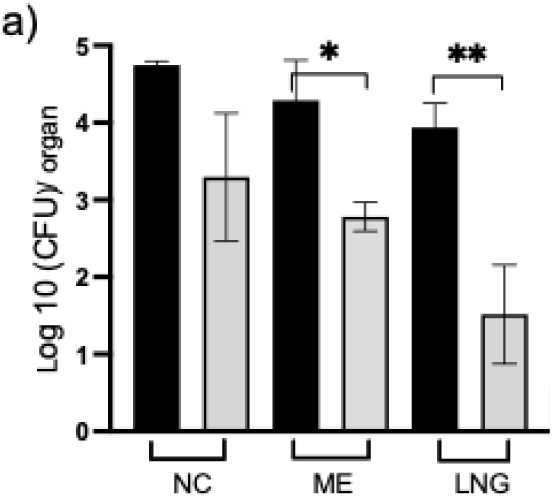
Effect of NPCC on *S. pneumoniae* colonization of adult mice. Graph shows CFU numbers of *S. pneumoniae* recovered from the nasal cavities (NC), Middle ears (ME) and lungs (LNG) of adult mice, 3 days after being challenged. The mice were either untreated (black columns) or mice treated with NPCC (grey columns). Error bars: S.E.M; *=p<0.05; **= p<0.001 (Results compiled from two experiments)

#### ii) B. pertussis

To examine the specificity of the effect of NPCC on the prominent gram-positive *S. pneumoniae*, we compared the relative protective effect of the NPCC on the gram negative respiratory and otopathogen *B. pertussis.* Groups of C57Bl/6J mice were either left untreated or inoculated with the NPCC, as above. Three days later both treated and untreated groups were intranasally challenged with ∼5 x10^5^ CFU of *B. pertussis* (gentamicin resistant) delivered in 10uL of PBS and colonization levels were assessed 3 days later. Figure 7 shows the combined results of two independent experiments. Untreated mice (-NPCC, black columns) had high numbers of *B. pertussis* in the NC (∼10^5^ CFU), as expected from the localized high dose intranasal challenge. The MEs also had relatively high but variable levels of the *B. pertussis* (∼ 3 to 5 x10^5^ CFU) indicating a robust ability to ascend the Eustachian tube to colonize and grow. Meanwhile the lungs of these mice showed relatively low numbers of pathogen (10s to 100s of CFUs), a result that we and others typically observe when delivering *B. pertussis* in low volumes initially restricted to the upper respiratory tract [51]. NPCC did not affect *B. pertussis* numbers in the NCs (∼5.5 x10^5^ CFU), but reduced the average in the MEs nearly 50-fold, with some variability, while the colonization of the LNGs appeared to show an increase in the colonization levels over the untreated group. These results, though not reflecting statistical significance (students t-test), suggest that the NPCC has a more modest impact on *B. pertussis* colonization of the MEs, suggesting a potentially more subtle effect of the NPCC on the progress of *B. pertussis* infections that may differ by organ in adult mice.

**Figure 7.**
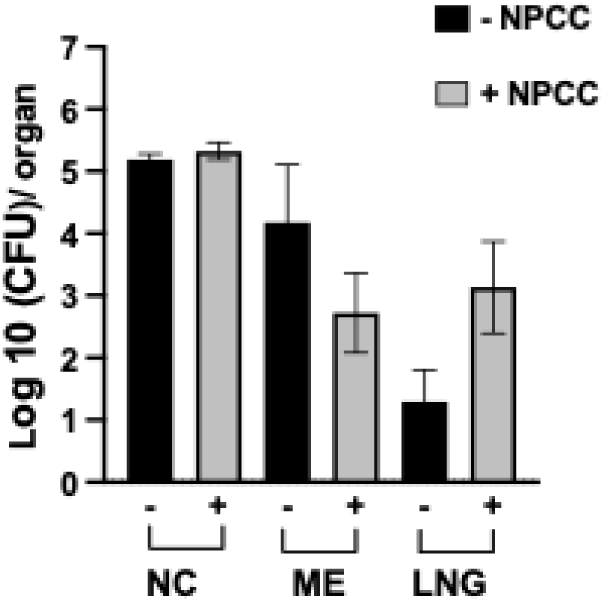
Effect of NPCC on *B. pertussis* colonization of adult mice. Graph shows CFU numbers of *B. pertussis* recovered from the nasal cavities (NC), Middle ears (ME) and lungs (LNG) of adult mice, 3 days after being challenged. The mice were either untreated (black columns) or mice treated with NPCC (grey columns). Error bars: S.E.M (Results compiled from two experiments).

### Transmission of NPCC

The natural establishment of the infant microbiota is strongly linked to the vertical transmission of maternal commensals to the infant [52, 53]. Here we examined whether dams inoculated with the NPCC would transmit these microbes to pups and compared this to how the NPCC would transmit among pups within a litter; expected to be facilitated by the close huddling behavior of nursing pups and their generally susceptibility to be colonized. For assessing vertical transmission, two days after giving birth (at P2, 10 pups) the female and male parent mice were inoculated with NPCC (∼10^5^ CFU in 10μL of PBS). To assess pup to pup transmission, 2 days after birth half the litter of pups (total 8 pups) were inoculated with the infant dose of the NPCC. The litters were then left undisturbed for three days to allow for any transmission following which the nasopharynx, middle ears and lungs of pups (now at P5) and parents were collected and culturable bacteria across the organs assessed. As indicated in Figure 8a, bacteria were recovered from the NCs of the two parent mice in relatively high numbers (∼5 x10^4^ CFU), but colonization levels for the pups remained low (∼10s to 100s of CFUs) across the nasopharynx, MEs and lungs of pups. These results suggest that, under these experimental conditions the inoculated parents do not appear to efficiently transmit the NPCC to pups. Further, in agreement with this observation,16S sequence analyses of the pup organ homogenates from the transmission experiments failed to detect any members of the NPCC (data not shown).

**Figure 8.**
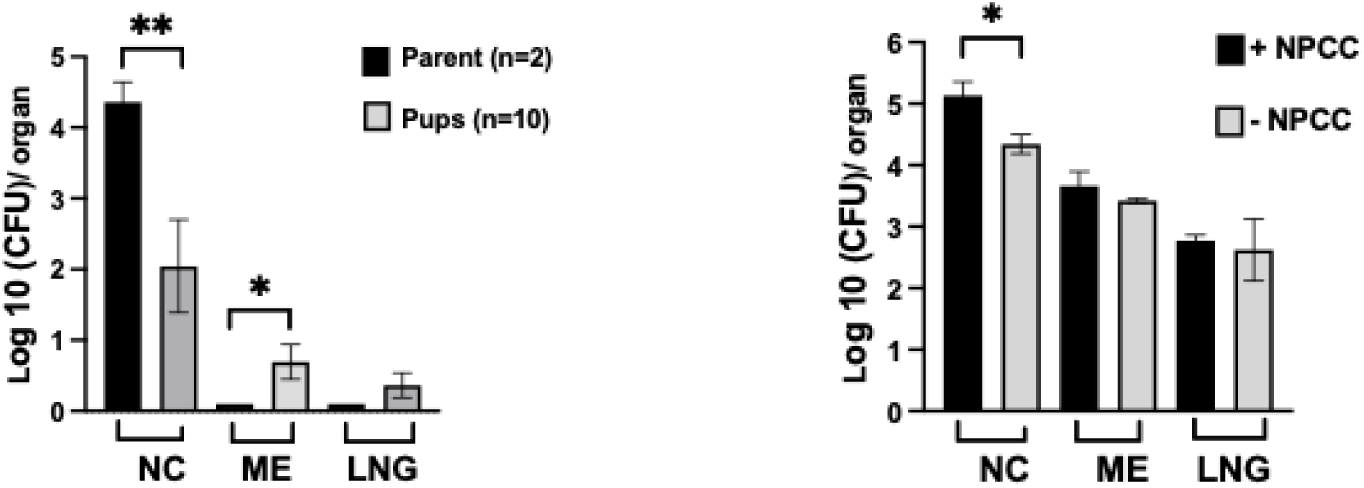
Transmission of the NPCC. a) *Parent to pup transmission*: Left graph shows CFU numbers of representative bacteria recovered from the nasal cavities (NC), Middle ears (ME) and lungs (LNG) of a single litter of C57BL/6J pups (grey columns), 3 days after the parent female and male mice were inoculated with NPCC (black columns). (Pup results compiled from 3 experiments) b) *Pup to pup transmission*: Graph shows CFU recovered from the nasal cavities (NC), Middle ears (ME) and lungs (LNG) of a single litter of C57BL/6J pups, 3 days after half the litter were either untreated (black columns) or mice inoculated with NPCC (grey columns). (Experiment done twice) Error bars: S.E.M; *=p<0.05; **= p<0.001.

In contrast to the differences in bacterial numbers seen across the organs for parent to pup transmission, very similar numbers of bacteria were recovered from both the inoculated and uninoculated pups within the same litter (Figure 8b) indicating that pup to pup transmission is highly efficient.

## DISCUSSION

Several studies have examined the respiratory (and middle ear) bacterial microbiota of patients suffering OM, comparing them with those of healthy (OM-free) controls to see if commensals might be contributing towards susceptibility or resistance to the disease [54]. The majority of 16S rRNA analysis of nasopharyngeal/adenoidal swab samples from OM patients have been found with expanded populations of either *S. pneumoniae*, *Moraxella catarrhalis* or *Hemophilus influenzae* often dominating their microbiota; in agreement with culture-based diagnostics of middle ear fluid identifying these as the major etiological agents of OM [55, 56]. However, a much greater variation of commensal species is observed among samples from healthy individuals making it difficult to identify a core protective microbiota [57]. In contrast to these case-control studies, the five health-associated commensals of the NPCC species we examine derives from the longitudinal analyses of a single cohort of infants suffering recurrent ear infections, reinforced by studies in similar cohorts. This has allowed us to directly link the change among commensal populations between “sickness” and “health”. Importantly, all the species selected for the NPCC used here have been associated with healthy controls in one or more of various studies [58–61].

Several salient features of our study need to be highlighted. The NPCC was identified in a human infant study and its protective effect was examined in infant and adult mouse models with two common pathogens that efficiently colonize mice but follow different infection schemes. We used an acute delivery model for the NPCC in neonates, and we accommodated the small size of the respiratory system by reducing the delivery volume to 2.5μL to initially localize bacteria to the nasal cavity. In addition, with the rapid day-to-day developmental changes of the infant immune system and relative transience of the infant microbiota [62], the dynamics of NPCC colonization for infant mice is expected to be different to that of adult mice.

NPCC treatment consistently resulted in a trend to reduce the bacterial load of both *S. pneumoniae* and *B. pertussis* in the MEs of both infants and adults. These results underscore a ME specific protective effect of the NPCC in agreement with its clinical healthy ME-associated status. Moreover, the ability of the NPCC to impede very severe, near lethal, pathogen challenges make this protection noteworthy. It should also be mentioned that the outcomes of neonatal mouse intranasal challenges vis-a vis ME infections are relatively uncharacterized and it is likely that modifications to the infant treatment/challenge scheme could enhance the protective effect of the NPCC. The relative changes in the population structures of *Limosilactobacillus reuteri* and *Lactobacillus johnsonni* in the lungs following NPCC administration points to how the benefits of protective commensals might also be mediated indirectly or orchestrated remotely through the expansion of other commensals at distal locations. There is the pressing question of whether the reduction in CFUs of the pathogens in the MEs of treated mice are attributable to a single species of the NPCC or from a symbiotic contribution of two or more species. A Finnish study singling out a specific commensal (*S. salivarius* [20]) as a probiotic failed to show OM protective effects [63], while other studies that intranasally delivered *Haemophilus haemolyticus* to mice [64], or a commensal *Pasteurella* species to humans [65] report beneficial effects on OM. In this respect, identifying the contributions of the individual NPCC members, while beyond the scope of the current study, will be important and is currently being planned.

Unlike for the MEs, pathogen colonization of the nasal cavities was little affected by NPCC in adults and pups alike. It is possible that the high dose pathogen challenge directly deposited in the NC overwhelmed the potential protective effect of the NPCC that might be observable in a more natural low dose challenge study. We and others have seen profound differences in colonization outcomes between high and low inoculum doses with bordetellae and *M. tuberculosis* [51, 66, 67] and in this respect it is possible that a challenge using a smaller dose to reflect a more natural exposure would reveal effects of the NPCC in the nasal cavity, generally the first site of pathogen colonization.

In the lungs, the effects of the NPCC on the spread and growth of the pathogen was only measurable for the adults, in which the inoculum was localized to the nasal cavities. NPCC treatment of the adults provided substantial protection (>100-fold lower numbers) against *S. pneumoniae* spreading to the lungs. In contrast to adults, the proportionally high volume (15μL) delivered to pups deposited the pathogens across the respiratory tract. In pups the NPCC was seen to provide substantial protection (>100-fold) against *S. pneumoniae*, but had no effect against *B. pertussis*; both NPCC-treated and untreated pups showed high lung colonization levels of *B. pertussis*. These observations underscore profound differences in how *B. pertussis* interacts with the NPCC compared to *S. pneumoniae* and point to a specificity to the impact that the resident microbiota have on the outcomes of different respiratory infections.

It was surprising that we detected no parent to pup transmission (experiment repeated 3 times) given what is generally understood of the susceptibility of the neonatal mucosal niche. However, we used a relatively short 3-day window of exposure to limit secondary pup-to pup transmission events which would be highly efficient due to the huddling behavior of pups. As such, high levels of shedding of bacteria such as *S. pneumoniae* and *B. pertussis* are detected in nasal swabs up to 7-10 days after inoculation. Additionally, microbiota transfer may occur most efficiently in the very earliest periods after birth.

Several factors can influence commensal health of the respiratory tract, the primary one being use of antibiotics. It is not uncommon for children with recurrent OM, to undergo multiple rounds of antibiotic treatment, compounding the risks of antibiotic resistance and further destabilizing commensal microbiota communities that might otherwise contribute to resisting pathogen invasion. Despite studies indicating only modest positive effects [68], parental demand for a solution, combined with the relative ease of prescribing antibiotics, makes OM the dominant reason for antibiotic prescription to children. Unfortunately, emerging evidence indicates antibiotics contribute to multiple prolonged or permanent effects on the microbiome with negative impacts on the immune and vaccine responses of treated infants [69, 70].

Potential paths out of the trap of repeated OM-antibiotics-OM cycles may be enabled by better understanding of how resident commensals resist/repel invading pathogens. Defining mechanisms of direct competition may reveal molecules that commensals secrete that are safe and effective to deliver within the respiratory tract. Understanding indirect competition may reveal key nutrients the targeted pathogen requires for growth or pathogenesis, allowing strategies to deplete those or rationally design antagonists to their uptake/utilization. Determining how commensals might induce host responses that minimally disrupt the healthy epithelia but effectively eliminate pathogens could allow those to be induced pharmaceutically or delivered topically. Whatever the mechanism involved, defining and understanding how commensals block pathogen invasion/pathogenesis has the potential to greatly reduce the great burden of OM, and the most common reason for the over use of antibiotics in children.

## Acknowledgements

Authors are grateful to the staff members of the Central Animal Facilities, University of Georgia for help in maintenance of experimental mice used in this study.

## SUPPLEMENTARY DATA

**Supplemental Figure 1.**
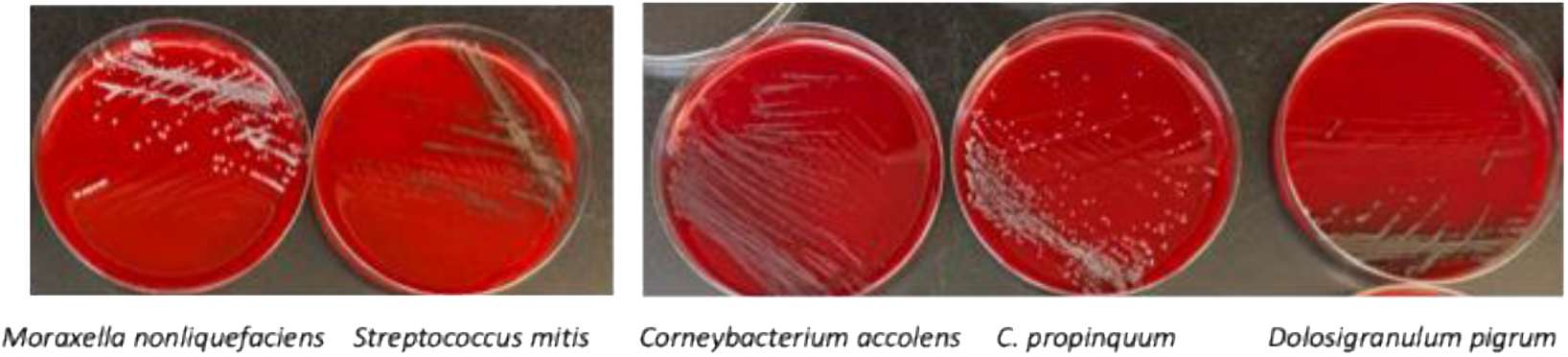
Individual NPCC strains grown at 37°C on Tryptic Soy agar supplemented with 5% defibrinated sheep blood. *M. nonliquefaciens* and *S. mitis* colonies detectable after 18 hours. *C. accolens*, *C. propinquum* and *D. pigrum* detectable after 48 hours

**Supplemental Figure 2.**
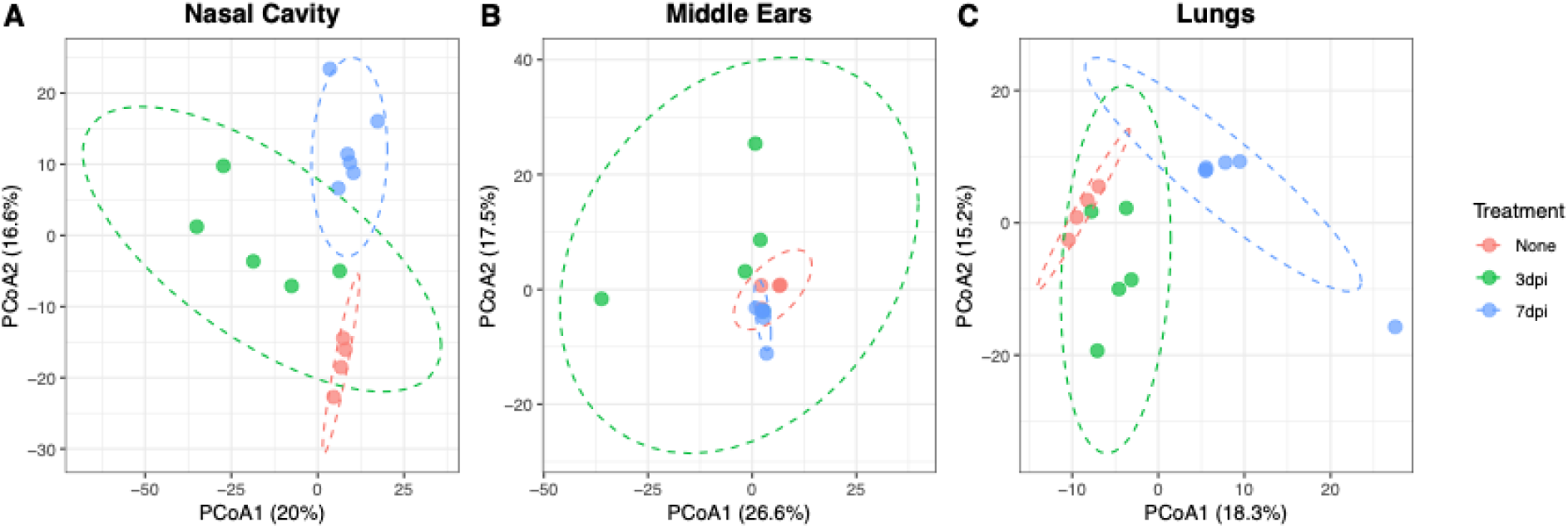
Impact of NPCC treatment on overall microbiome compositions of the respiratory tissues. Principle components analysis of nasal cavity (A), middle ear (B), and lung tissues (C) in untreated mice and in mice 3 and 7 days after inoculation of the NPCC.

